# Using Continuous Directed Evolution to Improve Enzymes for Plant Applications

**DOI:** 10.1101/2021.08.26.457776

**Authors:** Jorge D. García-García, Kristen Van Gelder, Jaya Joshi, Ulschan Bathe, Bryan J. Leong, Steven D. Bruner, Chang C. Liu, Andrew D. Hanson

**Author notes:** **Corresponding author:** Andrew D. Hanson. Author for communication (Andrew D. Hanson). Senior author (Andrew D. Hanson). **Author contributions:** A.D.H. conceived and designed the project with conceptual and practical guidance from C.C.L.; J.D.G.-G. performed experiments with support from K.V.G., J.J., U.B., and B.J.L.; S.D.B. carried out structural modeling; A.D.H. and J.D.G.-G. wrote the article with contributions from other authors. A.D.H. agrees to serve as author responsible for contact and ensures communication.

## Abstract

Continuous directed evolution of enzymes and other proteins in microbial hosts is capable of outperforming classical directed evolution by executing hypermutation and selection concurrently in vivo, at scale, with minimal manual input. Provided that a target enzyme’s activity can be coupled to growth of the host cells, the activity can be improved simply by selecting for growth. Like all directed evolution, the continuous version requires no prior mechanistic knowledge of the target. Continuous directed evolution is thus a powerful new way to modify plant or non-plant enzymes for use in plant metabolic research and engineering. Here, we first describe the basic features of the *Saccharomyces cerevisiae* OrthoRep system for continuous directed evolution and compare it briefly with other systems. We then give a step-by-step account of three ways in which OrthoRep can be deployed to evolve primary metabolic enzymes, using a THI4 thiazole synthase as an example and illustrating the mutational outcomes obtained. We close by outlining applications of OrthoRep that serve growing demands (i) to change the characteristics of plant enzymes destined for return to plants, and (ii) to adapt (‘plantize’) enzymes from prokaryotes – especially exotic prokaryotes – to function well in mild, plant-like conditions.

**One-sentence summary:** Continuous directed evolution using the yeast OrthoRep system is a powerful new way to improve enzymes for use in plant engineering as illustrated by ‘plantizing’ a bacterial thiamin synthesis enzyme..

## INTRODUCTION

### Continuous versus classical directed evolution

The directed evolution of enzymes is an iconic achievement of synthetic biology, marked by the award of the 2018 Nobel Prize in Chemistry to Frances Arnold (Arnold, 2019; Wang et al., 2021). Directed evolution as first conceived (classical directed evolution) involves iterative cycles of in vitro mutation at high rates (hypermutation) to diversify the target sequence, transformation of the resulting library of mutant sequences into cells (the platform), and screening or selection for the desired characteristics (Bershtein and Tawfik, 2008; Arnold, 2009; Engqvist and Rabe, 2019). Effective though classical directed evolution is, its reliance on separate diversification, transformation, and selection steps that are labor-intensive and lengthy severely restricts its throughput capacity (scale) and its access to fitness space requiring more than a very few mutations (depth) (Morrison et al., 2020; Rix and Liu, 2021).

Enter continuous directed evolution systems. After over a decade of development, these have recently come of age, and several are now available for *Escherichia coli* and yeast *Saccharomyces cerevisiae* hosts (aka ‘platforms’) that are well suited to evolving metabolic enzymes in vivo, e.g. OrthoRep (Ravikumar et al., 2018), EvolvR (Halperin et al., 2018), T7-DIVA (Álvarez et al., 2020), eMuta (Park and Kim, 2021), and CRAIDE (Jensen et al., 2021). Continuous systems overcome restrictions on depth and scale by operating entirely in vivo, targeting hypermutation specifically to the gene being evolved while simultaneously applying selection (Morrison et al., 2020; Rix and Liu, 2021). They all require the target gene’s function to be robustly coupled to a selectable phenotype. For metabolic enzymes, a standard way to do this is to use a platform strain that relies on the target enzyme for growth, e.g. to complement a loss-of-function mutation. Examples from the yeast OrthoRep system include evolving the malarial parasite dihydrofolate reductase for resistance to the antimalarial drug pyrimethamine in a *dfr1* disruptant (Ravikumar et al., 2018), and evolving the thermophilic *Thermotoga maritima* histidine synthesis enzyme HisA to function at moderate temperature in a *his6* deletant (Zhong et al., 2020).

Because continuous directed evolution has such potential in plant research and engineering, this article presents a user’s guide to a system – OrthoRep in yeast – that was one of the first enzyme-friendly systems developed (Ravikumar et al., 2014; Ravikumar et al., 2018) and with which we have the most experience (García-García et al., 2020). We begin below by describing OrthoRep and comparing it briefly with other systems mentioned above in the context of improving enzymes for use in plants.

### Features of the OrthoRep system

In OrthoRep (Ravikumar et al., 2018) (Figure 1), an engineered error-prone DNA polymerase (TP-DNAP1) specifically replicates a linear cytoplasmic plasmid (p1) that harbors the target gene, giving mutation rates as high as ∼10^−5^ substitutions per base replicated (s.p.b) without increasing the genomic mutation rate of ∼10^−10^ s.p.b. There are ∼10 copies of p1 per cell. The error-prone TP-DNAP1 is borne on a circular nuclear plasmid; it introduces mainly A:T→G:C transitions (∼70%) and G:C→A:T transitions (∼25%), with the four transversions at ≤1-2% each (Ravikumar et al., 2018).

**Figure 1.**
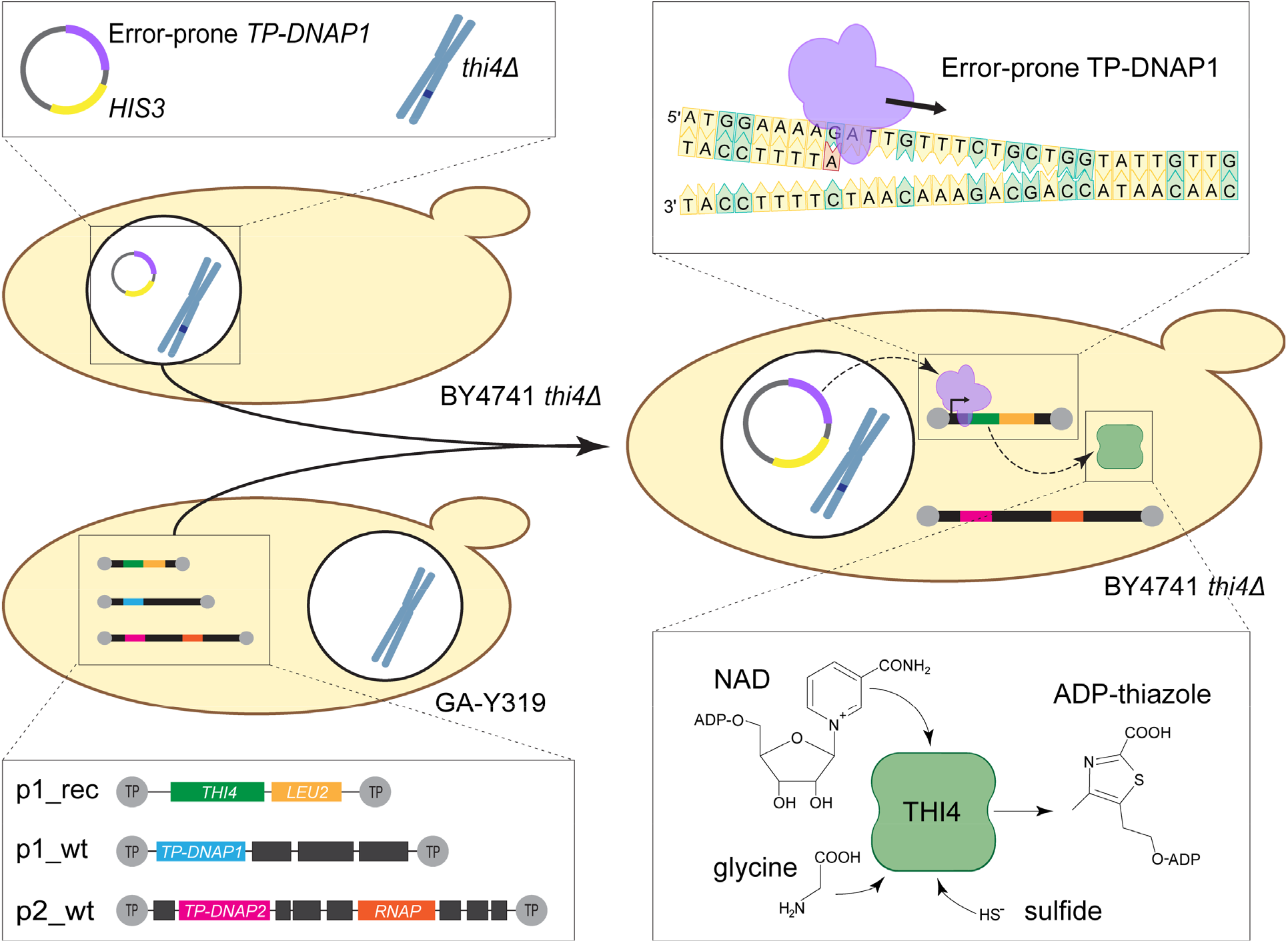
Overview of the OrthoRep system and its application to evolve prokaryote THI4 enzymes for improved performance in plant-like conditions. The OrthoRep mutagenesis machinery consists of three components: an orthogonal DNA polymerase, and two cytoplasmic plasmids (p1 and p2) originally from *Kluveromyces lactis*. The orthogonal DNA polymerase is an error-prone version of the terminal protein-primed DNA polymerase I (TP-DNAP1) of p1 that is transferred to a nuclear plasmid in a *thi4* knockout yeast strain (BY4741 *thi4*Δ). The *TP-DNAP1* of wild type p1 (p1_wt) is replaced by the target gene (*THI4*) and a *LEU2* selection marker via recombination, giving p1_rec. Protoplasts of strain GA-Y319 harboring the cytoplasmic plasmids and p1_rec are fused with the BY4741 *thi4*Δ strain containing error-prone *TP-DNAP1_611* to give BY4741 *thi4*Δ cells with p1_rec and wild type p2 (p2_wt). No selection pressure maintains p1_wt during or after protoplast fusion, so it is often not retained in the fusion strain. TP-DNAP1_611 replicates p1_rec and introduces mutations at a rate of ≥10^−6^ s.p.b. Mutated copies of p1_rec are transcribed by an RNA polymerase (RNAP) encoded on p2_wt. The THI4 target enzyme catalyzes synthesis of the thiamin precursor adenosine diphosphate-thiazole (ADP-thiazole) from NAD, glycine, and sulfide. Beneficial mutations in THI4 favor growth of the host cells in thiamin-free medium; deleterious mutations have the opposite effect.

A second linear cytoplasmic plasmid (p2) carries the components needed for transcription of genes on p1 as well as a DNA polymerase (TP-DNAP2) that replicates only p2. TP-DNAP1 and TP-DNAP2 use terminal proteins (TPs) covalently attached to the 5′ ends of p1 and p2, respectively, as replication origins. Because they are protein-primed, the p1 and p2 plasmids must be introduced into yeast cells via protoplast fusion, not regular transformation, and the target gene must be introduced into p1 using an integration vector and homologous recombination. Another special feature of OrthoRep is that the p1-encoded target gene transcripts are not polyadenylated by the host mRNA processing machinery and therefore benefit from the addition of a genetically encoded poly(A) tail of ∼75 nucleotides when higher expression is desired (Zhong et al., 2018). A <u>m</u>ulti<u>p</u>urpose p1 integration vector (GR-306MP) is available into which any target gene can be inserted between the 10B2 promoter and a poly(A) sequence in *E. coli* and then introduced into p1 by recombination in yeast (García-García et al., 2020).

To help appreciate OrthoRep’s mutational power, assume a mutation rate of 10^−5^ s.p.b., 10 copies of the p1-borne target gene per cell, and 2 × 10^7^ cells ml^-1^ at OD_600_ = 1 (Milo et al., 2010). It follows that, in a 3-ml culture grown from a starting OD_600_ of 0.05 to saturation (seven doublings, reaching an OD_600_ of 6.4), each base in the target gene is mutated a total of 3.8 × 10^4^ times in the population, i.e. a 1-kb target acquires mutations 38 million times in a single passage. Such high rates mean that single mutations at virtually all positions enter the population many times before any of them goes to fixation. Mutations that fix therefore tend to be those having the greatest selective advantage (Bailey et al., 2017) and the same advantageous mutations can fix frequently in independent populations (Ravikumar et al., 2018; Rix et al., 2020). Besides being very high, OrthoRep’s mutational power is durable, remaining completely stable over 90 generations, the maximum number explicitly tested (Ravikumar et al., 2018). A feature favoring durability is that in order to express the target gene on which the yeast cell has been made to rely, p1 must be replicated – and the error-prone TP-DNAP1 is the replicator. This kind of intrinsic selection for the error-prone propagation of the target gene is absent from some other continuous directed evolution systems, as discussed below.

Other advantages of OrthoRep are that it can be used in defined media (García-García et al., 2020), which is essential if the target gene complements a nutritional auxotrophy, and that expression of the target gene can be tuned (Zhong et al., 2018). Current disadvantages are that target gene expression cannot be turned on or off, and that the target gene’s promoter is mutated at the same high rate as its coding region (see below). An associated operational disadvantage of OrthoRep is that proving that a mutant sequence is responsible for the observed improvement in growth requires the exacting process of subcloning the evolved target gene from p1 and introducing it into fresh platform cells.

Lastly, cytoplasmic conditions in yeast, a eukaryote, are broadly at least as plant-like as *E. coli* in terms of pH, metabolites and cofactors, redox environment, and folding mechanisms (Kanehisa et al., 2006; Smith et al., 2007; Milo et al., 2010; Yébenes et al., 2011; Schwarzländer et al., 2016). OrthoRep is therefore well suited to evolve (‘plantize’) non-plant enzymes for future operation in plants.

### Other continuous directed evolution systems

EvolvR was first developed in *E. coli* and has since been implemented in yeast (Tou et al., 2020); T7-DIVA and eMuta were developed in *E. coli* and no other platforms are yet published. Table 1 compares EvolvR (Halperin et al., 2018), T7-DIVA (Álvarez et al., 2020), and eMuta (Park and Kim, 2021) with OrthoRep from a user’s point of view and we briefly expand below on five key points. The CRAIDE system for yeast (Jensen et al., 2021) is not covered as it has a far lower mutation rate than the others.

**Table 1.** User-guide comparison of continuous direction evolution systems in yeast and *E. coli*.

|  | OrthoRep | T7-DIVA | eMuta | EvolvR |
| --- | --- | --- | --- | --- |
| Mutation mechanism | Error-prone DNAP1 polymerase - plasmid pair | Base deaminase - T7 RNAP fusion | Base deaminase - T7 RNAP fusion | DNA polymerase III fused to CRISPR-guided nickase |
| Mutation rate (s.p.b)<sup>a</sup> | 10<sup>-8</sup>–10<sup>-5</sup> | Not determined | 9.4 × 10<sup>-5</sup> | 10<sup>-6</sup>–10<sup>-5</sup> |
| Mutation window | Up to at least 9 kb | Up to at least 951 bp | Up to at least 3 kb | ≤ ~60 bp |
| Minimal Media | Yes | Yes | Yes<sup>b</sup> | No |
| Plasmid or genomic insert | Plasmid | Genomic insert | Plasmid | Plasmid |
| Inducible target expression | No | Yes | Yes | Possibly |
| Mutational ON/OFF switch | No | Leaky | Yes | No |
| Promoter mutated | Yes | Optionally no | Yes | No |
| Tunability | Yes | No | No | No |
| Durability | TP-DNAP machinery tied to transcription of gene of interest in the cytoplasm | Gene expression does not depend on T7 RNAP, which will not maintain transcription | Gene expression depends on T7 RNAP, which maintains transcription | Mutagenesis decreases as the binding region of the sgRNA is mutated |
| Validation of mutants | Laborious | Laborious | Easy | Easy |

<sup>a</sup> cf. 10<sup>-9</sup> s.p.b. for wild type DNAP1 and spontaneous genomic mutation rates of ~10<sup>-10</sup> s.p.b. for yeast and ~2 × 10<sup>-10</sup> s.p.b. for *E. coli*.
<sup>b</sup> Determined in this study.

First, if truly continuous use is envisioned, i.e., many successive passages, the system’s mutational machinery must be stable, not prone to vitiation over time. As noted above, OrthoRep satisfies this condition since in OrthoRep, the target gene is unlikely to be expressed without being mutated because replication itself entails mutation. Similarly, eMuta satisfies this condition since the expression of the target gene uses the same T7 promoter that recruits the T7 RNA polymerase-based hypermutation machinery, with the caveat that mutations in the T7 promoter can change both the expression level of the target gene and its mutation rate. In contrast, EvolvR and T7-DIVA can potentially lose mutagenesis activity without losing expression: EvolvR due to mutations accumulating in the guide RNA binding sites and T7-DIVA because the base deaminase-T7 RNA polymerase fusion that makes mutations is independent of the endogenous RNA polymerase that drives expression, and imposes a fitness burden that predisposes it to loss (Kar and Ellington, 2018).

Second, in EvolvR, T7-DIVA, and eMuta the target gene’s promoter region is, or can be, largely insulated from mutation whereas in OrthoRep the promoter region cannot be insulated, opening the opportunity for promoter mutations to subvert the selection scheme simply by increasing target gene expression (see above). Such cheating could be minimized by using a promoter that is strong to begin with or eliminated by having a selection that cannot be satisfied by expression adaptation alone.

Third, EvolvR uses guide RNAs to direct the error-prone DNA polymerase to a narrow (∼60-bp) window within the target gene whereas OrthoRep, T7-DIVA, and eMuta systems mutate the whole target, working from one end to the other. To introduce mutations throughout the target, EvolvR therefore needs a series of guide RNAs, e.g., ∼16 for a 1-kb gene, and concatenating this many in the pEvolvR plasmid can be problematic. Without higher processivity error-prone DNA polymerases, EvolvR is more suitable for mutating a specific region or regions within a target gene using one or a few guide RNAs than it is for mutating the whole target.

Fourth, if evolution must be run in minimal or defined medium, as is common for metabolic selections, OrthoRep and T7-DIVA are suitable but EvolvR is unsuitable in *E. coli* as it has been found to require rich medium (García-García et al., 2020). eMuta is apparently suitable a priori and our preliminary tests support this, but published work so far has all been in rich medium.

Fifth, because the target gene in EvolvR and eMuta is on a regular plasmid it is straightforward to confirm that mutations in the target (not elsewhere) cause an observed improvement in growth as the evolved plasmid can be directly transformed into new cells. The integration of the target gene into the *E. coli* genome in T7-DIVA makes confirmation more laborious, and the location of the target on a protein-primed plasmid in OrthoRep does likewise, as noted above.

## RESULTS

### Background on the *MhTHI4* target gene used as an example

Yeast and canonical plant THI4s form the thiazole precursor of thiamin from NAD, glycine, and a sulfur atom (Chatterjee et al., 2011; Joshi et al., 2020). These THI4s use an active-site Cys residue as sulfur donor and irreversibly self-inactivate after a single reaction cycle, i.e., they are suicide enzymes. Degrading and resynthesizing suicidal THI4s is energetically inefficient, which makes plant THI4s targets for replacement with an energy-efficient, catalytic THI4 that mediates multiple reaction cycles (Hanson et al., 2018; Sun et al., 2019). Such catalytic TH4s, which use sulfide as sulfur donor, were first discovered in thermophilic, anaerobic archaea from hydrothermal vents, where sulfide levels are very high (millimolar) (Eser et al., 2016), and later found in mesophilic, anaerobic bacteria from other environments (Sun et al., 2019; Joshi et al., 2021). Prescreening of diverse catalytic THI4s by complementation of an *E. coli* thiazole auxotroph identified several with low but detectable activity in aerobic conditions. One of these, MhTHI4 from *Mucinivorans hirudinis* (an anaerobe from leech gut), was chosen for directed evolution in OrthoRep. The aim was to evolve (‘plantize’) MhTHI4 to operate more efficiently in air at a modest (plant-like) level of sulfide (Figure 1).

### Steps before deploying OrthoRep

First steps are to obtain and validate an appropriate yeast platform strain. In our case this was a *thi4* deletant of BY4741 (available from the yeast knockout YKO collection; Giaever and Nislow, 2014) in which *THI4* is replaced by a kanamycin/G418 resistance marker. Next steps are to codon-optimize the target gene for yeast and to remove targeting signals if present. An optional preparatory step we found useful is to check that high-level expression of the target gene from a regular plasmid enables at least some growth of the platform strain. For this check we used the CEN6/ARS4 nuclear plasmid ArEc-TDH3 carrying the *HIS3* marker and the constitutive TDH3 promoter to drive *MhTHI4* expression (Joshi et al., 2021). Gene expression driven by the TDH3 promoter on CEN/ARS plasmids is at least double that from the p1 plasmid, and in many instances much more than double (Zhong et al., 2018). Hence if a target gene does not pass this check, it is unlikely to support growth in the OrthoRep system, i.e., the activity of the target gene will fall below the selection window. (The selection window is the range of target gene activities within which selection can operate. In growth-coupled selection, the lower bound is detectable growth and the upper bound is when growth is no longer limited by target gene activity, i.e., when the strain being evolved grows as fast as wild type yeast in the same conditions.)

### Note on strains and selection media

To understand how the selection steps below work, note that strain GA-Y319, the source of wild type p1 and p2 plasmids (p1_wt and p2_wt), is auxotophic for uracil (*ura3*Δ*0*), leucine (*leu2*Δ*0*), histidine (*his3*), and tryptophan (*trp1*Δ*0*). Note also that the platform strain for selection in our case (as it would be in many others) was a BY4741 deletant from the YKO set that is auxotrophic for uracil (*ura3*Δ*0*), leucine (*leu2*Δ*0*), and histidine (*his3*Δ*1*) as well as thiamin (*thi4*Δ::KanMX). Thus GA-Y319 differs from BY4741 *thi4*Δ in requiring tryptophan, lacking kanamycin/G418 resistance, and not requiring thiamin.

### Construction of the strain to be evolved

This involves the operations described below and is summarized in Figure 2 as a flowchart.

**Figure 2.**
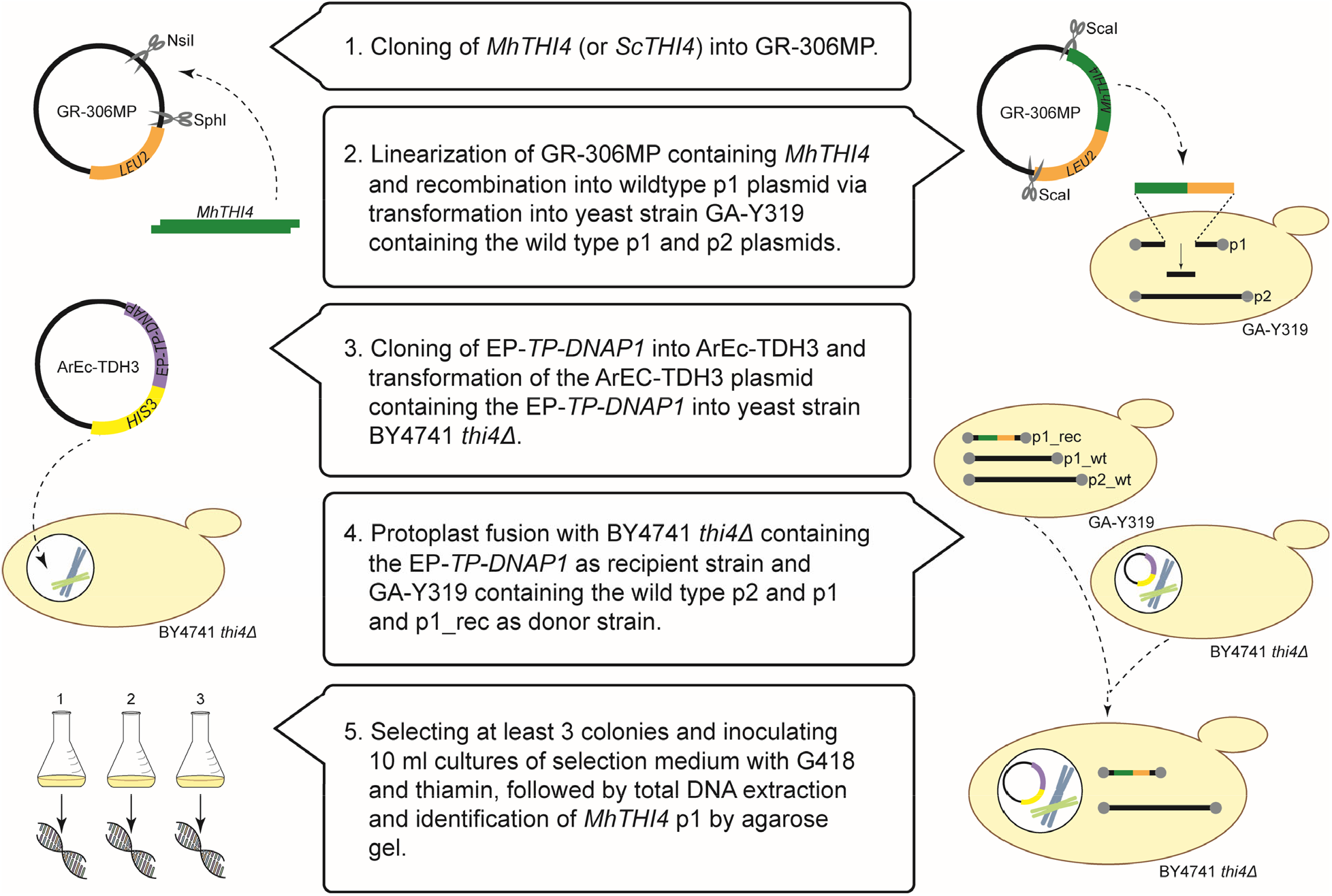
Flowchart of the construction of plasmids and strains involved in OrthoRep. Briefly, *MhTHI4* is cloned into the GR-306MP plasmid via restriction enzyme digest. The resulting construct is linearized for recombination into the p1_wt plasmid in strain GA-Y319. Additionally, strain BY4741 *thi4*Δ is transformed with the CEN/ARS plasmid containing an error-prone *TP-DNAP1* (*EP-TP-DNAP1*). A protoplast fusion between these two strains results in the BY4741 *thi4*Δ strain harboring p2_wt and p1_rec. Colonies from this fusion are cultured in selection medium with thiamin and total DNA is extracted and checked by agarose gel for the presence of p1_rec and p2_wt.

1. Clone the recoded target gene (and the corresponding yeast gene as positive control) in the multipurpose integration vector GR-306MP and prepare empty integration vector (as negative control).
2. Transform strain GA-Y319 carrying p1_wt and p2_wt with ScaI-digested GR-306MP integration vector containing the target gene and a *LEU2* marker, select colonies that grow in SC medium minus leucine, and screen whole-cell DNA by agarose gel analysis for the presence of the correctly recombined p1 plasmid (p1_rec) and p2_wt (Supplemental Figure S1). The p1_wt plasmid will also be present because at this stage it is the only source of the TP-DNAP1 DNA polymerase needed to replicate the recombinant p1 plasmid. The target gene in apparently correct p1_rec constructs is then PCR-amplified using whole-cell DNA as template and sequence-verified.
3. Transform the platform strain for selection (in our case, BY4741 *thi4*Δ) with the nuclear plasmid ArEc-TDH3 carrying the error-prone TP-DNAP1 polymerase and the *HIS3* marker, and select colonies that grow in SC medium minus histidine and tryptophan.
4. Carry out protoplast fusions between GA-Y319 carrying the recombinant p1_rec plasmid plus p2_wt (donor strain) and the platform strain (BY4741 *thi4*Δ) carrying the error-prone DNA polymerase (recipient strain), and select on SC medium minus leucine, histidine, and tryptophan (tryptophan is omitted to eliminate the GA-Y319 donor strain as noted above).
5. Pick colonies, culture in 10 ml of SC medium minus leucine, histidine, and tryptophan, plus G418 200 µg/ml, and in our case 300 nM thiamin. For this step, ammonium in the SC medium is replaced by glutamate because G418 is not compatible with ammonium. G418 is added to select for the *thi4* deletion instead of attempting to select for thiamin prototrophy at this stage, and to control bacterial contaminants. A 5-ml portion of the culture is taken for DNA isolation to check again by gel for the correct plasmid profile (Supplemental Figure 2A). If correct, the remaining 5 ml is the population to be evolved and is used to make an archival glycerol stock and for 3-ml precultures of SC with ammonium, minus leucine, histidine, and tryptophan, minus G418, plus thiamin 300 nM to start evolution experiments. Note that the p1_wt plasmid is usually visible in gels after protoplast fusion but disappears as evolution proceeds; other plasmid bands that may appear during evolution are generally inconsequential and can be disregarded (Supplemental Figure 2B).

### Implementing evolution strategies

Three basic strategies are shown in the flowchart of Figure 3. Even if the first strategy works, it is worth running the others as well because they can enable longer evolutionary paths to cross fitness valleys.

**Figure 3.**
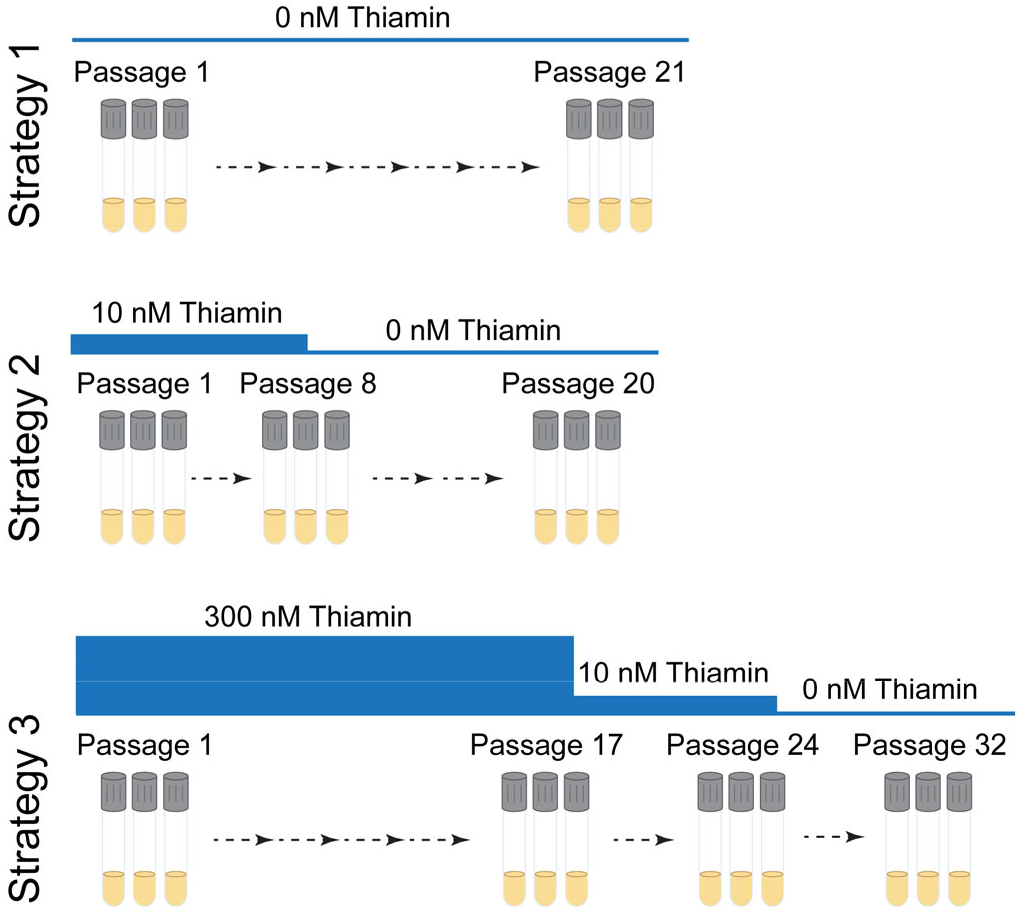
Flowchart of the three strategies implemented in OrthoRep. Strategy 1 was serial passaging in the absence of thiamin. Strategy 2 had two steps: eight passages in the presence of 10 nM thiamin, followed by 12 passages in absence of thiamin. Strategy 3 used stepwise decreases in thiamin supplementation, starting with 17 passages in 300 nM thiamin, followed by seven passages in 10 nM thiamin, and ending with eight passages in absence of thiamin.

1. Cold turkey: Selection is started as strictly as possible at once. In our case, this meant transferring well-washed cells straight to thiamin-free medium (initial OD_600_ = 0.05) and then serial passaging every six to ten days.
2. Gradual: Selective pressure is ramped up slowly, e.g., by tapering the supply of a required metabolite (thiamin in our case) or by ramping up the concentration of an inhibitor. We used a thiamin concentration (10 nM) low enough to limit growth to a final OD_600_ of ∼1, serially passaged cultures that grew to OD_600_ >1, then withheld thiamin completely.
3. Variant library building: Cells are cultured for many generations in nonselective conditions before starting selection. This allows neutral drift to generate a large library of mutant sequences on which selection can then act. We grew cells for ∼100 generations in the presence of a luxury thiamin concentration (300 nM), then moved successively to strategies 2 and 1. Note that this strategy can also be applied instead of in vitro mutagenesis to generate a mutant library for use in classical directed evolution (Wang et al., 2021).

As orientation on when to expect a beneficial mutation to detectably increase culture growth, consider this idealized case. Assuming a conservative doubling time of 20 h in minimal medium (García-García et al., 2020), if a mutation conferring normal growth rate arises in a non-growing population, the cell carrying this mutation will reach OD_600_ 1 in ∼22 days if there is no competition from other mutants. In practice, we stopped passaging populations that had no measurable growth after 12-20 days. Early termination is warranted for thiamin auxotrophs because thiamin is destroyed during use as an enzyme cofactor (Hanson et al., 2016); hence cells making no thiamin do not just stay in stationary phase but become thiamin-depleted and die. Other selection schemes would not have this special constraint.

All strategies require decisions about culture volumes and number of replicate populations to be used. Trade-offs of population size vs. number of independent populations are involved (Bailey et al., 2017; Rix et al., 2020; Rix and Liu, 2021): large populations increase the chance of rare mutational events, independent small populations stop different beneficial mutations competing with each other. Practically, we opted for 3-ml tube cultures and three independent populations per strategy for a small-scale exploratory operation. OrthoRep can be run on far larger scales in anything from large flasks to 96-well plates and up to 90 independent populations have been evolved in parallel (Ravikumar et al., 2018).

### Evaluating evolution outcomes

The growth of a population typically improves with successive passages until reaching a plateau (see Figure 4A for representative data); the population is then plated out to select, e.g., 5–10 single colonies. In the simplest case these colonies can go straight to sequencing of the p1-borne target gene and its promoter. However, we found it better to first eliminate cheaters that survive via cross-feeding from thiamin-producers by screening colonies for growth in thiamin-free medium. We revisit cheaters below.

**Figure 4.**
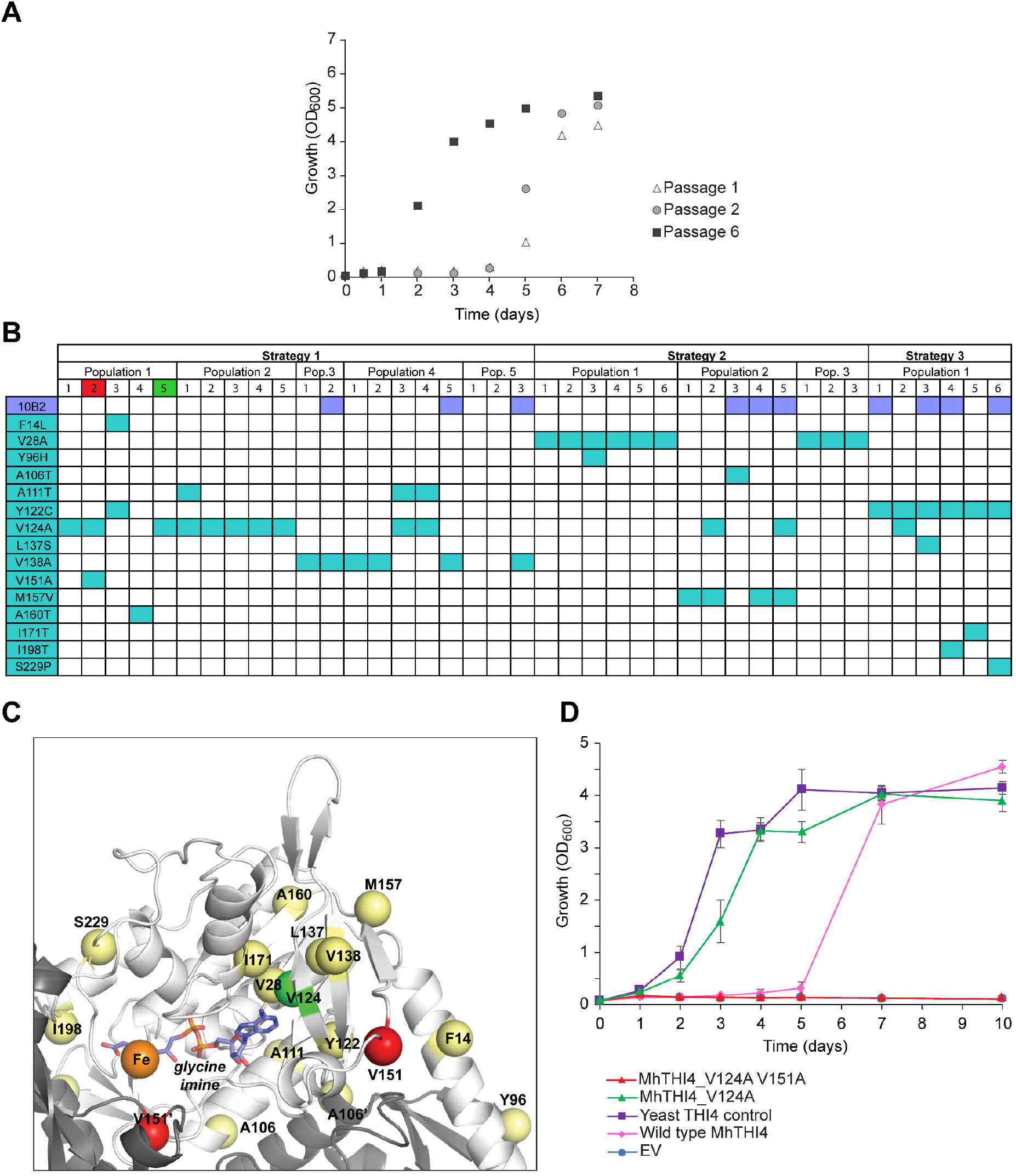
Representative outcomes for the evolution of the bacterial thiazole synthase MhTHI4. A, Growth improvement of a population harboring *MhTHI4* in response to selection in thiamin-free medium. Growth was substantially faster after six passages than after one or two passages. B, Table of mutations obtained in the promoter and coding regions. Fifteen nonsynonymous mutations were found in nine different populations evolved by the three strategies in Figure 3. Five mutations (V28A, A111T, Y122C, V124A, V138A) occurred in two or more independent populations. C, Structure model of MhTHI4 highlighting the 15 nonsynonymous mutations, which are shown in relation to the active site of one monomer of MhTHI4. Two positions where mutations have opposing effects, V124 and V151, are highlighted in green and red, respectively (also in part B); other mutations are highlighted in yellow. D, Populations harboring the single V124A mutant and double V124A V151A mutant were compared to the starting population harboring wild type *MhTHI4*, together with positive (yeast *THI4*) and negative (empty vector, EV) controls. Data are means ± SE of independent replicates (6 for *MhTHI4* V124A and V124A V151A, 3 for EV and yeast *THI4* controls, and 2 for the wild type *MhTHI4* check). The V124A mutation improved growth rate to near that of the yeast *THI4* control; adding the V151A mutation abolished growth.

Target genes that pass the above growth screen are then sequenced and genes with nonsynonymous mutations (Figure 4, B and C) are introduced into a fresh p1 plasmid in fresh platform cells, and the growth of the resulting strain is compared with that of the archived starting population to validate improvement (Figure 4D). Because quantitative in vitro assay of THI4 activity is not yet feasible, we could not confirm improved performance biochemically. This is somewhat unusual; reliable assays are available for most other enzymes so that improvements in mutant forms can be assessed directly in vitro.

## DISCUSSION

The exploratory work used here for demonstration is small-scale, but illustrates three key points about OrthoRep in particular, and continuous directed evolution in general. First is mutational power: from only nine independent populations (three per strategy), within weeks and without major manual interventions we recovered 15 nonsynonymous mutations, 17 synonymous ones, and 10 promoter mutations (Figure 4B and Supplemental Figure S3). Also, the same beneficial mutation (V124A) was recovered in five independent populations, and four more mutations (V28A, A111T, Y122C, V138A) were recovered at least twice. Such repeated evolution is likely only if mutation rates are high and population sizes are large (Bailey et al., 2017), and is prima facie evidence that the mutation is beneficial.

Because single mutations like V124A proved in the event to increase the activity of our target MhTHI4 enough to match that of native yeast THI4 (Figure 4), in retrospect it seems likely that such mutations could have been found by classical directed evolution methods. However, note that merely matching the activity of yeast THI4 is an extremely low bar because yeast THI4 is a suicide enzyme, i.e., mediates just one reaction cycle and dies (Chatterjee et al., 2011). That several apparently beneficial single mutations were recovered even in our small operation suggests that multi-mutational paths to higher activity very probably exist. As continuous directed evolution offers quick and seamless access to such paths, there is substantial scope for further use of OrthoRep to improve MhTHI4, as discussed next.

The second point is that the selection window (explained above) has a hard upper limit, reached when target enzyme activity evolves far enough to enable the yeast host to grow at a near-wild type rate, as with the V124A mutant (Figure 4D). Evolving the enzyme any further requires modification of the strategy. One option is to reduce the *amount* of the target enzyme by dialing down its expression (to make less of the enzyme work harder). OrthoRep is tunable and offers several ways to do this (Zhong et al., 2018). Another option is to reduce the in vivo *activity* of the target enzyme by adding a competitive inhibitor (e.g. a substrate analog) to the selection medium (Rix et al., 2020); for THI4 evolution the thiazole analog 2-aminohydroxyethylthiazole may be suitable (Iwashima et al., 1986). The next challenge is therefore to shift the selection window so as to unleash OrthoRep to find adaptive paths that involve multiple mutations and lead to higher activity than is achievable by single residue changes.

The third point is that evolving populations with high mutation rates are necessarily complex. One manifestation of the complexity is that, of 40 mutant genes sequenced, 27 had more than one mutation (some being synonymous and thus probably neutral). Multiple mutations in an evolving gene are obviously wanted because they enable longer evolutionary walks through fitness landscapes (Rix and Liu, 2021) but they also make it more difficult to pinpoint which mutations are causal. Another manifestation of population complexity is the appearance of the V151A mutation in a population improved by the beneficial V124A mutation even though V151A abolishes activity altogether (Figure 4D). The survival of this double mutant cheater can be explained by the ability of thiamin-producing yeast strains to secrete at least half their thiamin to the medium (Chatterjee et al., 2011) and thereby cross-feed non-producers (Haj-Ahmad et al., 1992). Although cheating via cross-feeding may be particularly favored for thiamin because thiamin is needed only in small amounts, cross-feeding can potentially weaken any selection in which growth is coupled to production of a metabolite.

The small dataset in Figure 4 also illustrates some general features of directed evolution, continuous or classical (Bershtein and Tawfik, 2008; Arnold, 2009). First, catalytic function can often be substantially improved by single residue changes, as in the case of the V124A mutation. This mutation, and the deleterious V151A mutation, illustrate a second feature: mutations with large effects are commonly in the active site, and rationalizable (Figure 4C). Thus, V124 is predicted to hydrogen bond with the glycine imine reaction intermediate and V151 is a conserved residue immediately adjacent to an invariant aspartate that binds the metal cofactor (Zhang et al., 2016). A third feature is that many mutations are distant from the active site, making the beneficial ones hard to rationalize.

## PERSPECTIVES

We close by highlighting two plant-related applications for OrthoRep (or other systems) for which there is a demand that is likely to grow, and by drawing a parallel with the recombinant DNA revolution.

The first application is changing the characteristics of an enzyme from a plant that is destined for return to the same plant via genome editing (Lin et al., 2020). This is an attractive crop improvement strategy because the US and many other countries exempt genome editing from GMO regulation of crop plants (Schmidt et al., 2020). The sequence of operations would be this: the plant, target enzyme, and desired change are defined, the selection scheme is designed, the enzyme gene (recoded if necessary) is evolved, the improved gene is sequenced, and the mutations that cause the improvement are edited into the native gene in the plant. Enzyme characteristics that could be modified this way include herbicide resistance (Dong et al., 2021), feedback inhibition (Alberstein et al., 2012), and in vivo working life (as measured by catalytic-cycles-till-replacement, CCR) (Hanson et al., 2021).

The second application is to adapt (‘plantize’) enzymes from prokaryotes – especially exotic prokaryotes – to function efficiently in mild, plant-like conditions. Our evolution of MhTHI4 is an actual case. The enzymes of the CETCH cycle (Schwander et al., 2016) and similarly daring metabolic designs (Bar-Even, 2018) are potential cases. The CETCH cycle is a synthetic pathway for CO_2_ fixation that is more efficient than natural plant pathways. However, it uses 17 different enzymes sourced from nine different organisms from three kingdoms of life, and has been implemented only in vitro using purified enzymes. As the enzyme sources include bacteria or archaea that come from near-anoxic environments and have different metabolic networks and metabolite profiles to plants, CETCH cycle enzymes are a priori likely to need plantizing before they work well in planta. Note that plantizing is conceptually akin to adapting enzymes to function in nonnatural environments, which is a standard application of classical directed evolution to improve enzymes for ex vivo industrial use (Arnold, 2009). It is also akin to directed evolution to adapt human enzymes for *E. coli* expression (Aharoni et al., 2004).

Lastly, continuous directed evolution is powerful, near-revolutionary, and so new that its applications are only just starting to be explored across the bioscience spectrum. Such exploration already extends beyond enzymes to engineering nucleic acid functions (Wang et al., 2021) and synthetic antibodies (Wellner et al., 2021). We can now extend this exploration into the plant world, much as happened a generation ago with recombinant DNA technology and its successful application in crop improvement.

## MATERIALS AND METHODS

### Plasmids and plasmid construction

Gene and vector sequences are given in Supplemental Table S1; primer sequences are in Supplemental Table S2. Media components are listed in Supplemental Table S3. The sequences of yeast *THI4* (*ScTHI4*) and yeast-codon optimized *Mucinivorans hirudinis THI4* (*MhTHI4*) without SphI, NsiI, EcoRI, or ScaI restriction sites were synthesized by GenScript® (Piscataway, New Jersey). Using Phusion™ High-Fidelity DNA Polymerase (Thermo Fisher Scientific, catalog no. F-530XL), *ScTHI4* and *MhTHI4* were PCR-amplified (total volume 25 µl) for cloning into ArEc-TDH3 or GR-306MP (García-García et al., 2020); the amplicons were gel-verified (2-µl aliquots were run on a 1% agarose gel) and column-purified (GeneJET PCR purification Kit, Thermo Fisher Scientific, catalog no. K0701). For cloning into ArEc-TDH3, 5 µg of empty vector and 1 µg of purified amplicon were digested overnight with 1 µl of SphI-HF® (New England Biolabs, catalog no. R3182) in a total volume of 50 µl. The mixtures were then heat-inactivated at 65°C for 20 min, and 5 µl of CutSmart buffer and 2 µl of EcoRI-HF® (New England Biolabs, catalog no. R3101) were added before incubation for 6 h at 37°C. The enzymes were again heat-inactivated and the digested DNA was column-purified. To build GR-306MP constructs, 10 µg of vector and 1 µg of amplicon were digested overnight with 2 µl (vector) or 0.5 µl (amplicon) each of SphI-HF® and NsiI-HF® (New England Biolabs, catalog no. R3127) in a total volume of 50 µl. After heat-inactivation at 65°C for 20 min, the digested GR-306MP backbone was separated from the *mKate2* fragment (García-García et al., 2020) by agarose gel; the backbone band was extracted using the GeneJET Gel Extraction Kit (Thermo Fisher Scientific, catalog no. K0691) and column-purified. The final plasmids were assembled in a total volume of 20 µl using 100 or 200 ng of linearized ArEc-TDH3 or GR-306MP backbone with molar insert:plasmid ratio of 7:1 (116 or 281 ng of digested PCR product, respectively) and 1 µl of T4 DNA ligase (Thermo Fisher Scientific, catalog no. EL0011). The mixtures were incubated for 3 h at room temperature; 2 µl was then used to transform *E. coli* TOP10 cells. Transformants were selected on LB plates containing 50 µg/ml kanamycin (ArEc-TDH3) or 100 µg/ml carbenicillin (GR-306MP). If no colonies appeared, the remaining ligation mix was incubated overnight at room temperature and transformed into TOP10. All constructs were sequence-verified. To avoid shortening of the poly(A) tail in GR-306MP constructs by repeated propagation, a fairly large batch of transformed TOP10 cells was grown in 50 ml of LB and a plasmid DNA stock was isolated with the Wizard® Plus Midipreps system (Promega, catalog no. A7640). The poly(A) tail length (73 bp) was verified by sequencing. The pAR-Ec611 (ArEc-Rev1-611) plasmid harboring medium-error-rate polymerase TP-DNAP1_611 (≥10^−6^ s.p.b.) was obtained from Addgene (Watertown, MA).

### Yeast strains

All media components for growing yeast are listed in Supplemental Table S3 and referred to below by their abbreviations as referenced in the table. The donor strain GA-Y319 (*MATa can1 his3 leu2*Δ*0 ura3*Δ*0 trp1*Δ*0 flo1* + p1_wt + p2_wt) (Ravikumar et al., 2014) was grown at 30°C in liquid YPD or on YPD plates. Strain BY4741 *thi4*Δ (*MATa his3*Δ*1 leu2*Δ*0 met15*Δ*0 ura3*Δ*0*; *thi4*Δ::KanMX) was obtained from Euroscarf (Oberursel, Germany) and grown at 30°C in liquid YPD + G418 or on YPD + G418 plates. To create the recipient strain for protoplast fusion, BY4741 *thi4*Δ was transformed with the error-prone polymerase in pAR-Ec611 (see above) as described (García-García et al., 2020).

### Complementation assay with the target gene in nuclear plasmid ArEc-TDH3

To transform BY4741 *thi4*Δ with ArEc-TDH3 harboring *MhTHI4*, the lithium acetate (LiAc) method (Mount et al., 1996) was used with modifications. A 3-ml YPD + G418 preculture was inoculated with BY4741 *thi4*Δ and grown to saturation at 30°C. Ten ml of YPD + G418 per transformation was inoculated with a 1:35 dilution of the preculture and grown to an OD_600_ of 0.6. Cells were pelleted by centrifugation (5 min, 1,500 *g*), washed with 10 ml of Milli-Q^®^ water per 10 ml of culture, recentrifuged, and washed with 300 µl of TE/LiAc (10 mM Tris-HCl, 1 mM Na_2_EDTA, pH 7.5, 100 mM LiAc) and then with 100 µl of TE/LiAc per 10 ml of culture. The washed pellet was resuspended in 267 µl of 50% PEG 3350 plus 40 µl of 10× TE (100 mM Tris-HCl, 10 mM Na_2_EDTA, pH 7.5) and 40 µl of 1 M LiAc and mixed with 25 µl of 5 mg/ml of single-stranded salmon sperm DNA and 300 ng of plasmid DNA (ArEc-TDH3, ArEc-TDH3_*ScTHI4*, or ArEc-TDH3_*MhTHI4*). After incubation for 45 min at 30°C with agitation, the cell suspension was gently mixed with 28 µl of DMSO and incubated for 20 min at 42°C. Cells were then pelleted by centrifugation (5 min, 700 *g*), resuspended in 1 ml of YPD and recovered for 1 h at 30°C and 220 rpm. Finally, the cells were washed with 1 ml of 0.9% NaCl, resuspended in 100 µl of 0.9% NaCl and plated on SC -His -Trp containing 300 nM thiamin; plates were then incubated for 3–5 d at 30°C. Single colonies were inoculated into 3 ml of SC -His - Trp containing 300 nM thiamin and grown for 2 d at 30°C and 220 rpm. After reaching stationary phase, 1 ml f the culture was harvested by centrifugation and washed five times with 1 ml of thiamin-free SC -His -Trp. The cells were then resuspended in 300 µl of the same medium and used to inoculate 3 ml of this medium at an OD_600_ of 0.05. Growth (OD_600_) was recorded after 12 h, 24 h, then every 24 h, and compared to the negative (ArEc-TDH3 empty vector) and positive (*ScTHI4* in ArEc-TDH3) controls (Joshi et al., 2021).

### Extraction of total DNA

Strains were grown for ∼2 d until saturation in 5 ml of appropriate selection medium. Cells were harvested by centrifugation (5 min, 1,500 *g*), washed with 1 ml of 0.9% NaCl, collected again by centrifugation, and resuspended in 0.5 ml of 1 M sorbitol, 0.1 M Na_2_EDTA, pH 7.5. To digest cell walls, the cell suspension was mixed with 5 µl of 5 U/μl Zymolyase (Zymo Research, catalog no. E1005) and incubated for 1 h at 37°C with agitation. Cells were then collected by centrifugation (5 min, 1,500 *g*) and resuspended in 0.5 ml of TE1 buffer (50 mM Tris-HCl, 20 mM Na_2_EDTA, pH 7.4). Fifty μl of 10% w/v SDS and 5 μl of 20 mg/ml proteinase K (Thermo Fisher Scientific, catalog no. EO0491) were mixed with the cell suspension and the mixture was incubated at 57°C for 30 min. After adding 200 μl of 5 M potassium acetate, the mixture was kept on ice for 1 h and then centrifuged (21,100 *g*, 5 min). Seven hundred µl of the supernatant were transferred to a 1.5-ml tube and mixed with 700 µl of isopropanol. The solution was mixed by inverting the tube and kept at room temperature for 10 min, then centrifuged (21,100 *g*, 5 min); the nucleic acid pellet was dried for 15 min at room temperature, resuspended in 300 μl of TE2 buffer (10 mM Tris-HCl, 1 mM Na_2_EDTA, pH 7.4) and 1.5 µl of 10 mg/ml RNase (Thermo Fisher Scientific, catalog no. EN0531) was added, followed by incubation at 37°C for 30 min. After adding 30 μl of 3 M sodium acetate and 200 µl of isopropanol, the solution was mixed by inverting the tube and held for 10 min at room temperature. The tube was then centrifuged (21,100 *g*, 5 min). The pellet was dried as above and resuspended in 50 μl of TE2. After centrifugation (21,100 *g*, 10 min), the supernatant containing total DNA was transferred to a new 1.5-ml tube. Successful isolation of p1 and p2 plasmids was verified by running 1 µg of total DNA on a 0.8% agarose gel. Note that detection of p1 plasmids can require overexposure during image capture due to their low abundance.

### Construction and validation of recombinant p1 plasmid

Ten µg of plasmid GR-306MP_*MhTHI4* were digested overnight using 1.5 µl of ScaI-HF® (New England Biolabs, catalog no. R3122) in a total volume of 50 µl, then heat-inactivated for 20 min at 65°C. A 2-µl aliquot was run on a 1% agarose gel to check for complete digestion, and the remaining 48 µl was used to transform strain GA-Y319 as above. Transformants were selected on SC -Leu for 5 d at 30°C. To confirm recombination of *MhTHI4* into p1, single colonies were inoculated into 5 ml SC -Leu and grown to saturation for ∼2 d at 30°C and 220 rpm. Total DNA was isolated as above and 1 µg was run on a 0.8% agarose gel to verify the presence of p1_wt, p1_rec, and p2_wt. The presence of the 10B2 promoter, *MhTHI4*, and the poly(A) tail was checked by PCR using total DNA as template and primers p1_F and Ribozyme_R (Supplemental Table S2) and sequencing of the amplicons.

### Protoplast fusion

Donor strain GA-Y319 harboring p1_rec and recipient strain BY4741 *thi4*Δ harboring the error-prone polymerase were grown in 3 ml of appropriate selection medium to saturation (∼2 d) at 30°C and 220 rpm. Cultures were diluted 1:50 into 50 ml of YPD and incubated overnight at 30°C. An aliquot of culture was transferred to a 50-ml tube and centrifuged (3,000 *g*, 5 min) to retain 0.3–0.4 g wet weight of cells (at OD_600_ of 4.5, 20 ml of culture gave ∼0.45 g of cells). Pellets were gently resuspended in 10 ml of Milli-Q^®^ water, centrifuged as above, and weighed to confirm recovery of 0.3–0.4 g of cells. Pellets were gently resuspended in 0.2% v/v β-mercaptoethanol and 0.06 M Na_2_EDTA to a final volume of 1.8 ml by tapping the side of the tubes. The cells were then incubated at 30°C without agitation for 30 min (inverting several times at 15 min), then collected by centrifugation as above, washed with 3 ml 0.6 M KCl with gentle resuspension, and recentrifuged. The pellets were resuspended in 4.8 ml buffer I (13.6 mM citric acid, 50.9 mM Na_2_HPO_4_, pH 6.1, 0.6 M KCl, 10 mM Na_2_EDTA) with 6 U/ml Zymolyase and incubated at 30°C for 1 h with occasional gentle inverting, after which the suspensions were centrifuged (700 *g*, 10 min, 0°C). The pellets were washed twice with 3 ml of 0.6 M KCl, resuspended gently and centrifuged as above with final resuspension in 3 ml of buffer I. At this point, the donor and recipient strains were mixed 1:1 (final volume 3 ml) by gently inverting the tube. The mixture was centrifuged (700 *g*, 10 min, 0°C), resuspended in 5 ml of buffer II (33% w/v PEG 3350, 0.6 M KCl, 50 mM CaCl_2_) and incubated at 30°C for 30 min with occasional inverting. The mixture was centrifuged as above and resuspended in 5 ml of buffer III (0.6 M KCl, 50 mM CaCl_2_). The cell mixture (250–500 µl) was added to 25 ml of SC -His -Leu -Trp +KCl +agar, warmed to 42°C, mixed gently by inverting and poured into a petri dish. Three colonies from each protoplast fusion were selected after 5-7 d and inoculated into 10 ml of SC -His -Leu -Trp +G418 and grown to saturation (2-3 d). Five ml of this culture (the ‘parental strain’) was harvested for total DNA extraction as above and run on an 0.8% agarose gel to confirm the presence of p1_rec and p2_wt. The remaining 5 ml was used to make a glycerol stock and to inoculate 3-ml cultures at a 1:100 dilution for gene evolution trials.

### Gene evolution trials

Strain BY4741 *thi4*Δ harboring p1_rec and p2_wt was cultured in 3 ml SC -His -Leu -Trp, without or with thiamin depending on selection strategy. All strategies were run at 30°C with 220 rpm shaking.

#### Strategy 1 (cold turkey)

3-ml cultures in SC -His -Leu -Trp with 300 nM thiamin were inoculated at a 1:100 dilution from the parental strain (see above) and grown to saturation (2–3 d). One ml of the culture was centrifuged (1,500 *g*, 2 min). The pellets were washed five times with thiamin-free SC - His -Leu -Trp, resuspended in 300 µl of this medium and used to start (at OD_600_ = 0.05) new 3-ml thiamin-free SC -His -Leu -Trp cultures. Cultures were grown until OD_600_ reached 3–5 (typically 4–6 d), then subcultured at a 1:100 dilution (starting OD_600_ 0.04–0.05) into fresh selection medium. OD_600_ was measured every 24 h for six passages and thereafter only before each new passage. Cultures that did not grow after 12-20 days were discarded. Cultures were analyzed for mutations after 6 (population 1), 14 (populations 2 and 3), or 21 (populations 4 and 5) passages (Figure 4B).

#### Strategy 2 (gradual)

3-ml cultures were grown and washed as in Strategy 1, used to start (at OD_600_ = 0.05) 3-ml cultures in SC -His -Leu -Trp containing 10 nM thiamin, and grown until OD_600_ was 1–5 for eight passages. Cells were then subcultured into thiamin-free SC -His -Leu -Trp, grown to OD_600_ of 3–5 for 12 passages, and analyzed for mutations.

#### Strategy 3 (variant library-building)

3-ml cultures of the parental strain were used to inoculate (1:100 dilution) 3-ml cultures of SC -His -Leu -Trp containing 300 nM thiamin. Cultures were grown in this medium to OD_600_ of 3–5 for 17 passages (∼100 generations). Cells were then transferred to SC –His -Leu -Trp plus 10 nM thiamin (gradual strategy) and grown to OD_600_ 3–5 for seven passages, followed by eight passages in thiamin-free SC -His -Leu -Trp. Cultures were then analyzed for mutations.

### Evaluation of mutant THI4s

Cultures to be analyzed for mutations were plated on SC -His -Leu -Trp +G418 with 10 nM thiamin and incubated at 30°C for 3 d. Selected colonies from plates were used to inoculate a 3-ml culture in thiamin-free SC -His -Leu -Trp and grown to saturation. A 10-ml culture in thiamin-free SC -His –Leu -Trp was inoculated with cells from the 3-ml culture at a 1:20 dilution and grown for at least 3 d. Five to seven ml were taken for total DNA extraction and the rest was used to make a glycerol stock. The *MhTHI4* gene was amplified from total DNA using primers outside the promoter, target gene, and poly(A) tail (primers p1_F and Ribozyme_R, Supplemental Table S2) and sequenced.

Selected mutant *MhTHI4* genes were re-cloned into GR-306MP, transformed into the GA-Y319 strain and fused with the BY4741 *thi4*Δ strain as above. At least three colonies from the protoplast fusion were each used to inoculate a 10-ml culture in SC -His -Leu -Trp +G418 and grown to saturation. Five ml of culture was harvested for DNA extraction as above and run on an 0.8% agarose gel to identify p1_rec and p2_wt. The other 5 ml was used to inoculate 3-ml cultures (1:100 dilution) in SC -His -Leu -Trp containing 300 nM thiamin and grown to saturation. Three 3-ml cultures in the same selection medium were inoculated from three colonies each of the empty vector control, the yeast THI4 positive control and the parental strain (MhTHI4 BY4741 *thi4*Δ yeast before evolution trials) and grown to saturation. One ml of the saturated cultures was centrifuged (1,500 *g*, 2 min) and washed five times with thiamin-free SC -His -Leu -Trp. The pellets were resuspended in 300 µl of this medium and the OD_600_ was taken. New 3-ml cultures in the same medium were inoculated from the resuspended pellets (at OD_600_ = 0.05) and grown for 10 d, recording OD_600_ on days 1–5, 7, and 10.

### Homology modeling of the MhTHI4 structure

The MhTHI4 homology model was based on the *Methanocaldococcus jannaschii* THI4 structure (PDB code 4Y4M) and sequence alignment. The figure was prepared with PyMol.

## Supplemental Data

The following materials are available in the online version of this article.

**Supplemental Figure S1**. Agarose gel of total DNA extracted from individual colonies of GA-Y319 transformed with the ScaI-digested GR-306MP plasmid.

**Supplemental Figure S2**. Expected agarose gel band patterning of total DNA extracts of BY4741 *thi4*Δ cells harboring *MhTHI4*.

**Supplemental Figure S3**. Mutations found in the promoter and coding regions of *MhTHI4*.

**Supplemental Table S1**. Sequences of vectors and genes used in this study. **Supplemental Table S2**. Primers used to clone and sequence *ScTHI4* and *MhTHI4*. **Supplemental Table S3**. Recipes of media used in OrthoRep.

## Acknowledgments

We thank the late Dr. Dan Tawfik for insightful advice about directed enzyme evolution.

